# Epigenetic characterization of pseudogenes across human tissues

**DOI:** 10.1101/2025.10.05.680540

**Authors:** Yunzhe Jiang, Beatrice Borsari, Mark Gerstein

## Abstract

Pseudogenes have long been considered non-functional remnants of genome evolution. However, compared to other noncoding elements, their epigenetic regulation remains poorly understood, largely due to their sequence similarity to parent genes and the lack of systematic functional analyses. Here, we leverage matched transcriptomic and epigenomic data from the ENCODE EN-TEx project, spanning ∼30 human tissues, to comprehensively profile pseudogenes and compare them to protein-coding transcripts and lncRNAs. Our analyses reveal epigenetic differences among pseudogene biotypes. Even when transcribed, processed pseudogenes—unlike protein-coding transcripts and unprocessed pseudogenes—lack canonically active histone marks and open chromatin signatures at their promoters. Instead, these regions are frequently associated with retrotransposon sequences, enriched for YY1-binding sites, and often engage distal enhancer-like elements through long-range genomic contacts. These findings suggest that processed pseudogenes may adopt regulatory mechanisms akin to transposable elements such as LINEs, relying on YY1-mediated enhancer-promoter looping or other distal interactions to facilitate transcription. By publicly releasing our catalog of pseudogene promoter regions, we provide a comprehensive resource for investigating the epigenetic and functional roles of pseudogenes in the human genome.

## Introduction

Since the release of the first draft in 2001, considerable efforts have been devoted to improving the annotation of the human genome^1–4^. For decades, most studies have focused on the comprehensive characterization of protein-coding genes, which constitute roughly 2% of the human genome, while largely neglecting the significance of noncoding elements, primarily long noncoding RNAs (lncRNAs), microRNAs (miRNAs), and pseudogenes. More recently, the advent of high-throughput sequencing technologies has made it possible to annotate, quantify, and epigenetically characterize the non-coding portion of the genome, particularly lncRNAs and miRNAs^5–7^.

On the other hand, pseudogenes have received much less attention and remain poorly characterized as they have long been considered non-functional remnants, often labeled as “junk DNA.” Pseudogenes are segments of DNA that resemble functional protein-coding genes but lose coding ability because of genetic disablements, such as the emergence of premature stop codons or frameshift mutations^8^. Protein-coding genes that give rise to pseudogenes are referred to as parent genes, and they are functional paralogs of pseudogenes. Thus, studying pseudogenes can be technically challenging due to their sequence similarity to parent genes, requiring accurate annotation of the human genome^9^.

Pseudogenes are classified into three biotypes based on their creation mechanisms^10^ (**Figure 1A**): unprocessed pseudogenes, which derive from the duplication of functional protein-coding genes; processed pseudogenes, which derive from the retrotransposition of mature messenger RNAs (mRNAs); and unitary pseudogenes, which accumulate fixed disablements in a given species (e.g., humans) while retaining functional orthologs in others (e.g., mice or primates). Roughly 10% of pseudogenes in the human genome are transcribed in a tissue-specific manner, with expression levels comparable to those of protein-coding genes^11^. Some pseudogenes also regulate the expression of their parent genes^12–15^. Given these regulatory roles, it is essential to understand their transcriptomic patterns and epigenetic regulation across tissues and cell types and how this regulation differs from that of protein-coding genes and lncRNAs.

**Figure 1.**
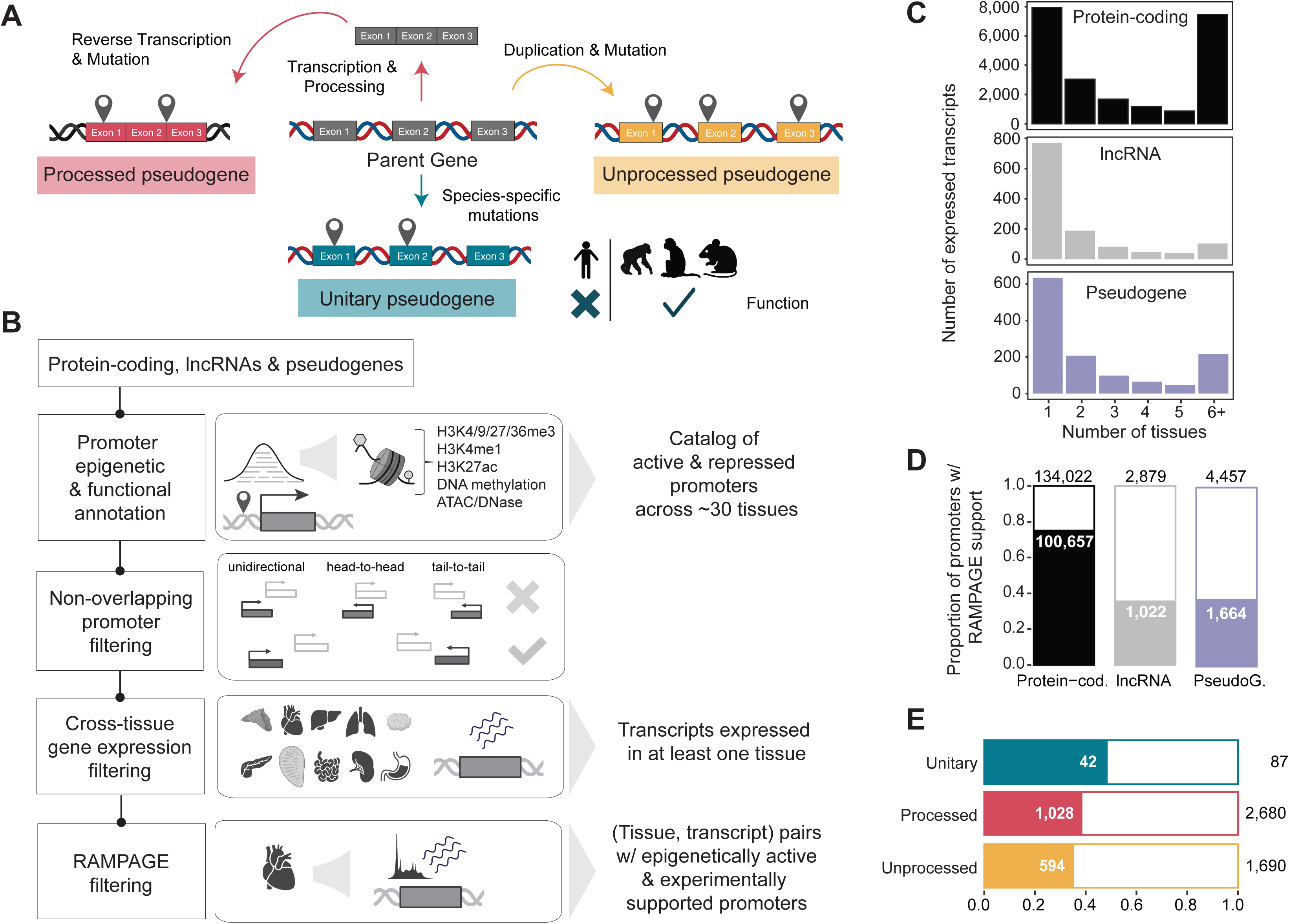
Building a catalog of epigenetically and functionally annotated promoters in the human genome. **(A)** Schematic representation of different pseudogene biotypes based on their mechanisms of origin, depicted in distinct colors. *(Left)* Processed pseudogenes, which arise from retrotransposition events. *(Middle)* Unitary pseudogenes, which result from mutations in functional protein-coding genes leading to a fixed loss of function in a given species. *(Right)* Unprocessed pseudogenes, which originate from gene duplication followed by functional decay. **(B)** Overview of the study design workflow. A catalog of active and repressed promoters was constructed across ∼30 human tissues using tissue-specific epigenome data. Transcripts were subsequently filtered based on three criteria: *(i)* annotation, ensuring no overlapping promoters; *(ii)* expression, requiring an average TPM ≥ 1 across donors and replicates; and *(iii)* RAMPAGE support for promoters. **(C)** Barplot showing, for different gene biotypes, the number of expressed transcripts (*y*-axis) across an increasing number of tissues (*x*-axis). These numbers are computed upon applying the cross-tissue gene expression filtering step in **(B)**. **(D)** Proportion of promoters, categorized by gene biotypes, that have RAMPAGE support, counting each transcript-tissue pair separately. **(E)** Analogous representation to **(D)**, but categorized by three pseudogene biotypes. “Protein-cod.”: protein-coding; “PseudoG.”: pseudogene. Panels **(A)** and **(B)** were created with <u>Biorender.com</u>.

In this study, we leveraged the EN-TEx^11^ multi-tissue transcriptomic and epigenomic dataset to comprehensively characterize pseudogenes across nearly 30 human tissues. We first annotated the epigenetic status of protein-coding, lncRNA, and pseudogene promoters, creating a publicly available resource for exploring the epigenetic and functional landscape of pseudogenes. Mining this dataset, we identified distinct epigenetic patterns not only between protein-coding and pseudogene transcripts but also among different pseudogene biotypes. In particular, our findings show that even when expressed, processed pseudogenes lack canonical active histone marks at their promoters. Instead, their promoters are strongly associated with retrotransposons and enriched for binding sites of the multifunctional transcription factor Yin Yang 1 (YY1). These results highlight the unique regulatory strategies of processed pseudogenes and provide new insights into their functional roles in the human genome.

## Results

### Building a multi-tissue catalog of epigenetically and functionally annotated promoters for pseudogenes, lncRNAs, and protein-coding transcripts

With the goal of characterizing the epigenetic landscape of pseudogenes and comparing it to other gene biotypes, we first compiled a comprehensive catalog of promoter regions for pseudogenes, lncRNAs, and protein-coding transcripts annotated with their epigenetic activity (**Figure 1B**; **Table S1**). To achieve this, we leveraged the EN-TEx epigenomic maps, including Chromatin Immunoprecipitation Sequencing (ChIP-Seq) data for six histone modifications (H3K27ac, H3K4me3, H3K27me3, H3K36me3, H3K4me1, and H3K9me3), as well as DNase I hypersensitive site sequencing (DNase-Seq), Assay for Transposase-Accessible Chromatin using Sequencing (ATAC-Seq), and DNA methylation profiling (see Methods). This resource offers a uniform characterization of promoter epigenetic activity across 26 human tissues. Using this, we defined promoters as ± 1000 base pair (bp) windows around the transcription start sites (TSSs) of GENCODE v29 transcripts. Then we annotated their activity based on the presence of active and repressive epigenetic features (see Methods). In addition to this epigenetic characterization, we also added to the catalog the functional properties of these promoters by analyzing their GC content, evolutionary conservation, and mutation spectra based on single nucleotide variant (SNV) data from the Genome Aggregation Database (gnomAD)^16^ (**Table S2**).

We then used this catalog to investigate the relationship between chromatin state and isoform expression across different gene biotypes. To minimize potential biases from overlapping transcripts of different genes, we first curated a subset of non-overlapping promoter regions (**Figure 1B**; see Methods). Next, we integrated matched RNA Sequencing (RNA-Seq) data at the isoform level from the same 26 human tissues (**Figure 1B**; see Methods). Specifically, for each gene in a given tissue, we selected the most highly expressed transcript, requiring an average expression level of ≥ 1 transcript per million (TPM) across all four EN-TEx donors (**Figure 1C and Figure S1**). Of these transcripts, we finally retained those whose promoters were experimentally supported by the presence of RNA Annotation and Mapping of Promoters for the Analysis of Gene Expression (RAMPAGE) peaks^17^ (**Figure 1B**). This analysis highlighted that promoters of protein-coding transcripts have stronger experimental support (75.1%) compared to those of lncRNAs (35.5%; Fisher’s exact test, odds ratio = 5.48, *p*-value < 2.2x10^-16^) and pseudogenes (37.3%; Fisher’s exact test, odds ratio = 5.06, *p*-value < 2.2x10^-16^) (**Figure 1D**). On the other hand, we did not observe strong differences in the proportion of RAMPAGE-supported promoters across different pseudogenes biotypes (**Figure 1E**). Together, we have uniquely included 335 pseudogenes, 490 lncRNAs, and 14,315 protein-coding transcripts across all tissues. We examined transcript-tissue pairs to investigate the interplay between transcript expression and histone modifications at the level of individual tissues for subsequent analyses.

### Processed pseudogenes lack canonically active histone marks at their promoters despite being expressed

As expected^18^, most protein-coding promoters exhibit active chromatin marks in the same tissue in which they are expressed, particularly H3K27ac and H3K4me3, with each covering over 75% of the total promoter length. In contrast, the extent of chromatin accessibility is lower, with peaks from ATAC-Seq and DNase-Seq covering an average of 42.4% and 23.8% of the promoter length, respectively (**Figure 2A**). lncRNA promoters exhibit intermediate levels of active mark coverage (∼60%), while pseudogene promoters have the lowest (∼25%) (**Figure 2A**). The lack of active chromatin signatures at pseudogene promoters raised the question of whether this pattern is a general feature of all pseudogenes or specific to a particular biotype. Upon closer examination, we found that promoters of unprocessed and unitary pseudogenes exhibit epigenetic profiles more similar to those of lncRNAs and protein-coding transcripts, while processed pseudogenes are significantly depleted of active histone marks (**Figure 2B**).

**Figure 2.**
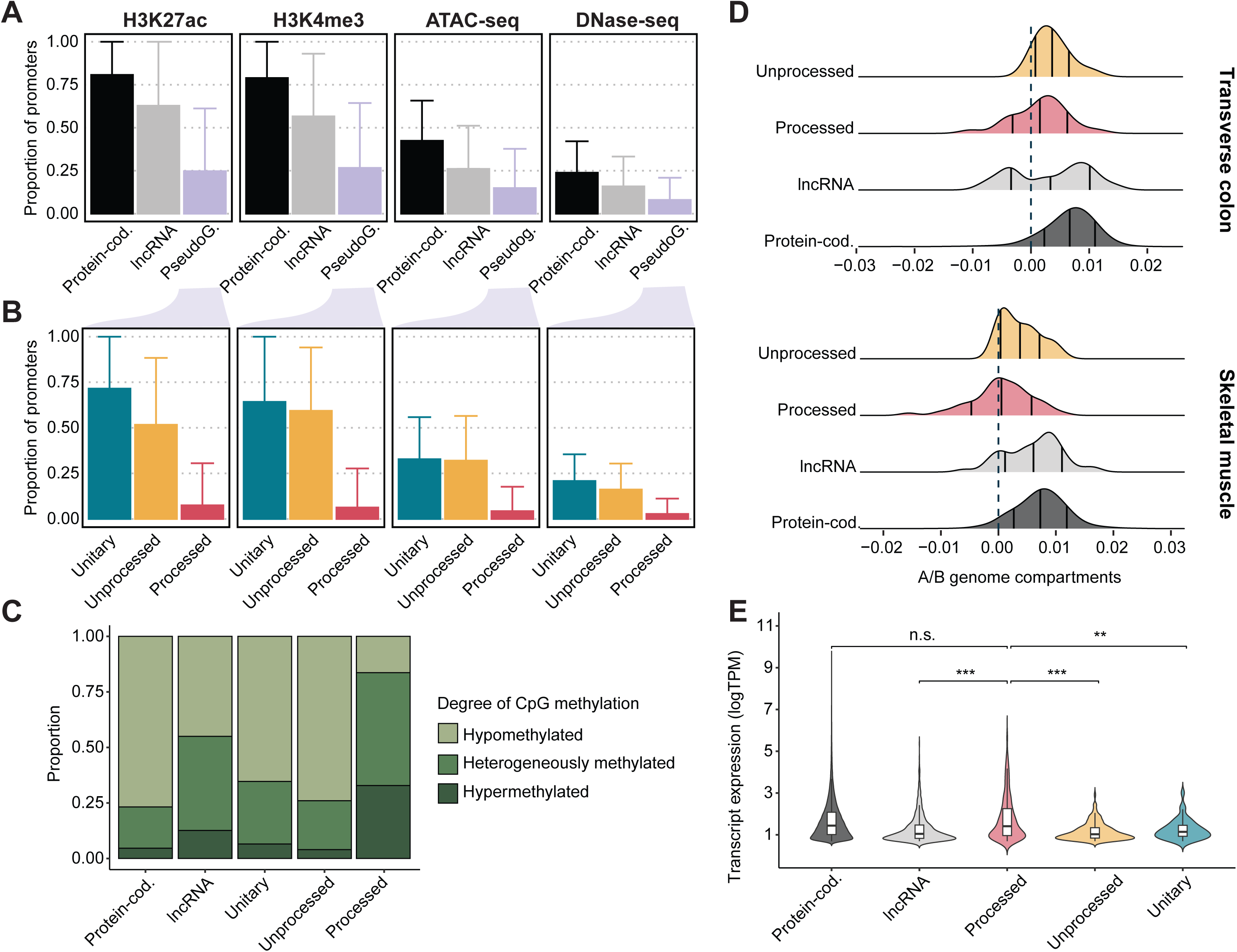
Epigenetic characterization of transcribed promoters. **(A)** Average proportion of promoter bases (*y*-axis) of protein-coding, lncRNA, and pseudogene transcripts (*x*-axis) that overlap peaks of H3K27ac, H3K4me3, ATAC-Seq, and DNase-Seq. Proportions were calculated for each transcript in each tissue and then aggregated across tissues for each gene biotype. Error bars represent standard deviations across tissues. **(B)** The average proportion of bases in promoters for three pseudogene biotypes, calculated and aggregated as in **(A)**. Error bars represent standard deviations across tissues. **(C)** Percent stacked bar plot showing, for each gene biotype, the proportion of promoter CpG sites with different levels of methylation across tissues. **(D)** Ridgeline plot showing, for different gene biotypes, the distribution of transcript promoters found within active (A) and inactive (B) genome compartments in transverse colon *(Top)* and skeletal muscle *(Bottom)*. Unitary pseudogenes were excluded due to their small number of expressed transcripts in these tissues. Vertical lines indicate the mean and one standard deviation above and below the mean. **(E)** Violin plot showing the expression levels of transcripts from different gene biotypes. Processed pseudogenes were used as the reference group for the two-sided Wilcoxon rank-sum test. ***, *p*-value < 0.001; **, *p*-value < 0.01; n.s., not significant. “Protein-cod.”: protein-coding; “PseudoG.”: pseudogene.

These results align with the distribution of repressive chromatin features, particularly DNA methylation (see Methods). When categorizing promoters based on the number of methylated CpG sites in their matched expression tissue, we found that processed pseudogenes exhibit higher DNA methylation coverage. That is, they are more frequently heterogeneously methylated or hypermethylated compared to unprocessed and unitary pseudogenes, lncRNAs, and protein-coding transcripts (**Figure 2C and Figure S2**). Thus, they appear to be not only depleted of active chromatin features but also enriched in repressive signals. Furthermore, analysis of *in situ* Hi-C data available for the transverse colon and skeletal muscle revealed that processed pseudogene promoters are more frequently located in inactive genomic compartments (e.g., B compartments) (see Methods) (**Figure 2D**).

To rule out the possibility that these epigenetic differences are driven by underlying differences in expression, we compared the distribution of expression levels across tissues and gene biotypes. We found no significant difference in expression between processed pseudogenes and protein-coding transcripts (Wilcoxon rank sum test, *p*-value = 0.954, FDR-adjusted) (**Figure 2E**). Notably, despite their lack of active histone marks and open chromatin signatures, processed pseudogenes are even more expressed than unprocessed pseudogenes (Wilcoxon rank sum test, *p*-value = 1.98x10^-31^, FDR-adjusted).

### Sequence features reveal similarities between processed pseudogenes and retrotransposons

Our findings so far indicate that, across multiple tissues, processed pseudogenes lack features of active chromatin at their promoter, despite exhibiting similar or even higher expression levels than protein-coding transcripts or unprocessed pseudogenes. Hence, we explored sequence features to further investigate molecular mechanisms underlying these distinct regulatory patterns. Since processed pseudogenes originate from the retrotranscription of spliced mRNAs and typically lack intronic regions (**Figure 1A**), analyzing promoter sequences downstream of the TSS could introduce biases across gene biotypes. These regions differentially overlap exons in processed pseudogenes and introns in protein-coding transcripts, lncRNAs, and unprocessed and unitary pseudogenes, which are known to exhibit distinct conservation patterns^19^ and mutation rates^20^. To mitigate this bias, we focused specifically on the 1000-bp region upstream of the TSS.

First, we observed that these regions in processed pseudogenes have significantly lower GC content compared to other gene biotypes (Wilcoxon rank sum test, *p*-value = 1.04x10^-27^ vs. protein-coding transcripts, and *p*-value = 4.42x10^-19^ *vs.* unprocessed pseudogenes, FDR-adjusted) (**Figure 3A and Figure S3**). Considering that high GC content is a hallmark of promoters^21^, this finding is overall consistent with our results on the differential enrichment of active and repressive chromatin features.

**Figure 3.**
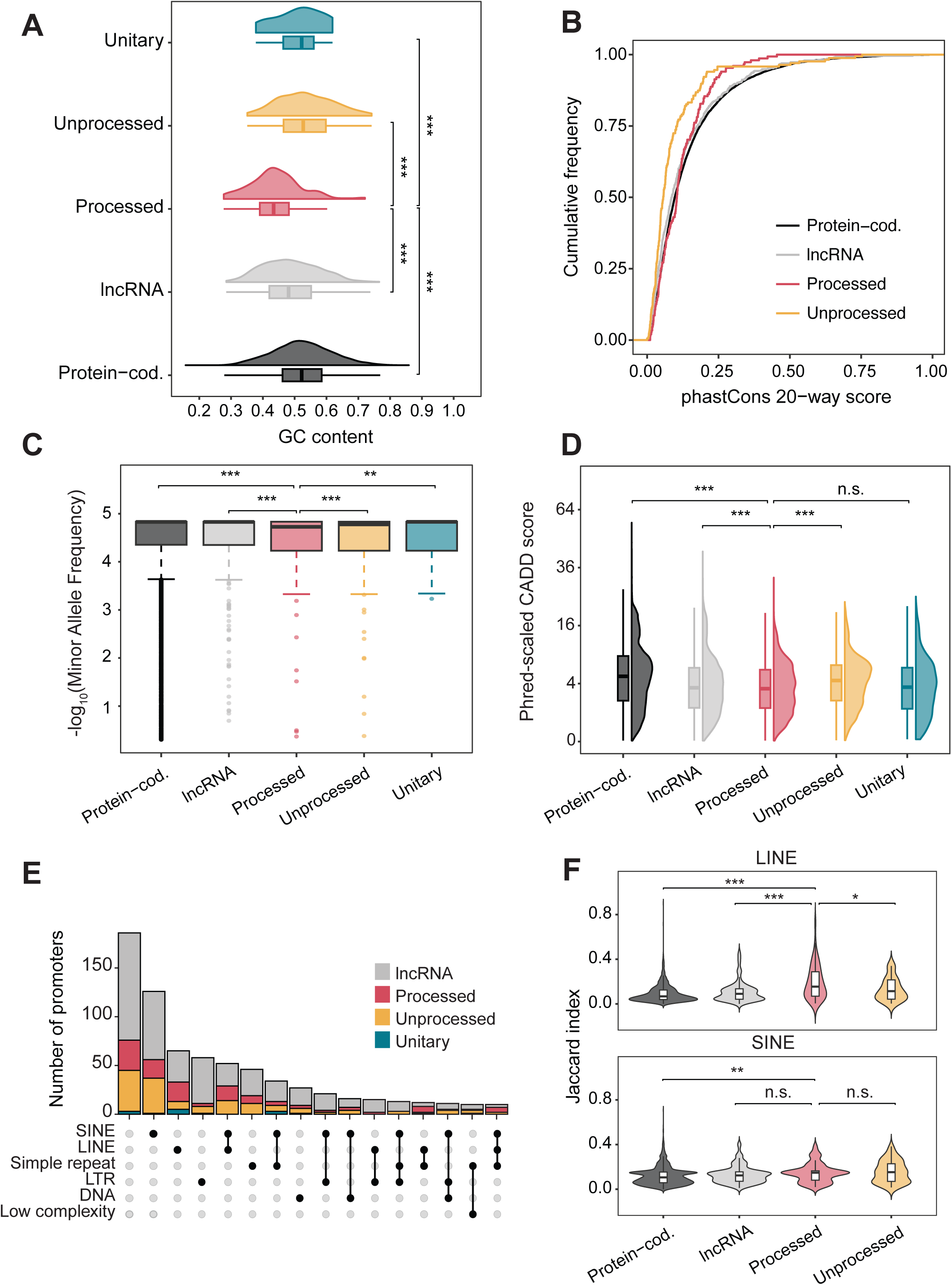
Sequence features of promoter regions. **(A)** Raincloud plots show the GC content of upstream regions of TSS for transcripts from different gene biotypes. Processed pseudogenes were used as the reference group for the two-sided Wilcoxon rank sum test. ***, *p*-value < 0.001. **(B)** Cumulative distribution of conservation scores (20-way phastCons) for regions upstream the TSS across different gene biotypes. Unitary pseudogenes were excluded due to the small sample size (N = 17). **(C)** Box plots displaying the distribution of MAF of SNVs within the upstream regions of TSS for different gene biotypes. Processed pseudogenes were used as the reference group for the two-sided Wilcoxon rank sum test. Due to the large sample size, 0.5% of outlier SNVs were randomly selected for visualization. Processed pseudogenes were used as the reference group for the two-sided Wilcoxon rank sum test. ***, *p*-value < 0.001; **, *p*-value < 0.01. **(D)** Raincloud plots depicting the distribution of phred-scaled CADD scores for SNVs within the upstream regions of TSS across different gene biotypes. Due to the large sample size, only 0.5% of SNVs classified as outliers were randomly selected for visualization. Processed pseudogenes were used as the reference group for the two-sided Wilcoxon rank sum test. ***, *p*-value < 0.001; n.s., not significant. **(E)** Upset plot showing the number of lncRNA and pseudogene promoters overlapping with various transposable elements. Some intersection groups were excluded due to the small size of elements (N < 10). **(F)** Violin plots display the degree of similarity (measured by the Jaccard index) between promoters of different gene biotypes and LINEs (*Top*) and SINEs (*Bottom*). Processed pseudogenes were used as the reference group for the two-sided Wilcoxon rank sum test. ***, *p*-value < 0.001; **, *p*-value < 0.01; *\*, p*-value < 0.05; n.s., not significant. “Protein-cod.”: protein-coding.

Next, we assessed inter- and intra-species conservation across promoter sets by analyzing the distribution of their average phastCons scores and the minor allele frequencies (MAF) of their single nucleotide variants (SNVs), respectively (see Methods). Our key finding is that upstream regions of TSS in processed pseudogenes exhibit a conservation profile higher than those in unprocessed pseudogenes, especially over long evolutionary timescales, and the pattern closely resembles that of protein-coding transcripts, particularly around the mode. (Wilcoxon rank sum test, *p*-value = 0.541 vs. protein-coding transcripts, and *p*-value = 2.02x10^-5^ vs. unprocessed pseudogenes; FDR-adjusted) (**Figure 3B and Figure S4**). This might suggest some degree of selective pressure acting on these fixed inter-species differences. However, this selection signal is less evident within the human population, as processed pseudogenes show an allele frequency distribution shifted towards more common variants (Wilcoxon rank sum test, *p*-valu*e* = 2.63x10^-21^ vs. protein-coding transcripts, and *p*-value = 1.31x10^-8^ vs. unprocessed pseudogenes, FDR-adjusted) (**Figure 3C and Figure S5A**). On the other hand, the proportion of ultra-rare variants in upstream regions of TSS in processed pseudogenes (82.1%) is, on average, lower than those in protein-coding transcripts (84.8%) but higher than those in unprocessed pseudogenes (80.6%) (**Figure S6**). This pattern is also consistent with their overall lower Combined Annotation Dependent Depletion (CADD) scores (**Figure 3D and Figure S5B**), suggesting that they tend to accumulate less impactful variants (Wilcoxon rank sum test, *p*-value = 0 vs. protein-coding transcripts, and *p*-value = 1.48x10^-137^ vs. unprocessed pseudogenes, FDR-adjusted).

Epigenetically, the promoters of processed pseudogenes show strong similarities with repetitive elements in the human genome^22^. In fact, most transposable elements (TEs) are enriched in DNA methylation^23^ and H3K9me3^24^, similar to the promoters of processed pseudogenes (**Figure S7**). However, despite this epigenetic silencing, some TEs, especially the long and short interspersed nuclear elements (LINEs and SINEs), are known to be transcriptionally active under certain conditions^25–27^.

Thus, we explored the possibility that there could be some shared regulatory mechanisms between TEs and processed pseudogenes. We analyzed the overlap between the different sets of promoters and annotated TEs (**Figure 3E and Figure S8**; see Methods). Compared to protein-coding transcripts, we observed that a substantial portion (39.3%) of promoters of processed pseudogenes overlap with LINEs, whereas SINEs are more enriched in the promoters of unprocessed pseudogenes (50.3%). Furthermore, the degree of sequence overlap (as measured by the Jaccard index) with both LINEs and SINEs is significantly different among different biotypes (Kruskal-Wallis test, *p*-value = 7.74×10⁻ for LINEs and *p*-value = 2.14×10⁻ for SINEs) (**Figure 3F**). Again, processed and unprocessed pseudogenes showed different patterns, reporting the highest overlap with LINEs and SINEs, respectively.

### YY1-binding sites are enriched at the promoters of processed pseudogenes and potentially mediate their interaction with distal regulatory elements

The strong genetic and epigenetic similarity between processed pseudogenes and LINEs prompted us to investigate whether they might share similar regulatory mechanisms. LINEs typically possess an internal RNA polymerase II promoter, enabling autonomous transcription, whereas SINEs depend on LINEs for transcription and mobilization^28^. Based on this, we hypothesized that the expression of processed pseudogenes could be mediated by a similar mechanism. To this end, we employed Homer2^29^ to identify transcription factor (TF) motifs enriched at the promoters of processed pseudogenes that could potentially mediate their transcription. Our analysis revealed a significant enrichment of the YY1 TF motif in ∼10% of promoter regions of processed pseudogenes, compared to the ∼1% in those of protein-coding transcripts and unprocessed pseudogenes (**Figure 4A**). A prior study had demonstrated that the 5’-untranslated region (5’-UTR) of human LINE-1 harbors a consensus YY1-binding site, which is a component of the LINE-1 core promoter and essential for accurate transcription initiation^30^. More recently, functional YY1-binding sites have been identified in the 5’-UTR monomers of the Tf_I subfamily of LINE-1 in mouse, where YY1 acts as a transcriptional activator, paralleling its role in human LINE-1^31^. We also found an enrichment of the YY2 motif in ∼30% of promoter regions of processed pseudogenes, compared to the ∼10% in those of unprocessed pseudogenes (**Figure 4B**). As the human paralog of YY1, YY2 shares the same consensus sequence binding as YY1 but with much lower affinity, while adopting a more ordered conformational state^32–34^. Together, these findings support our hypothesis that processed pseudogenes and human LINE-1 share similar regulatory mechanisms underlying their expression.

**Figure 4.**
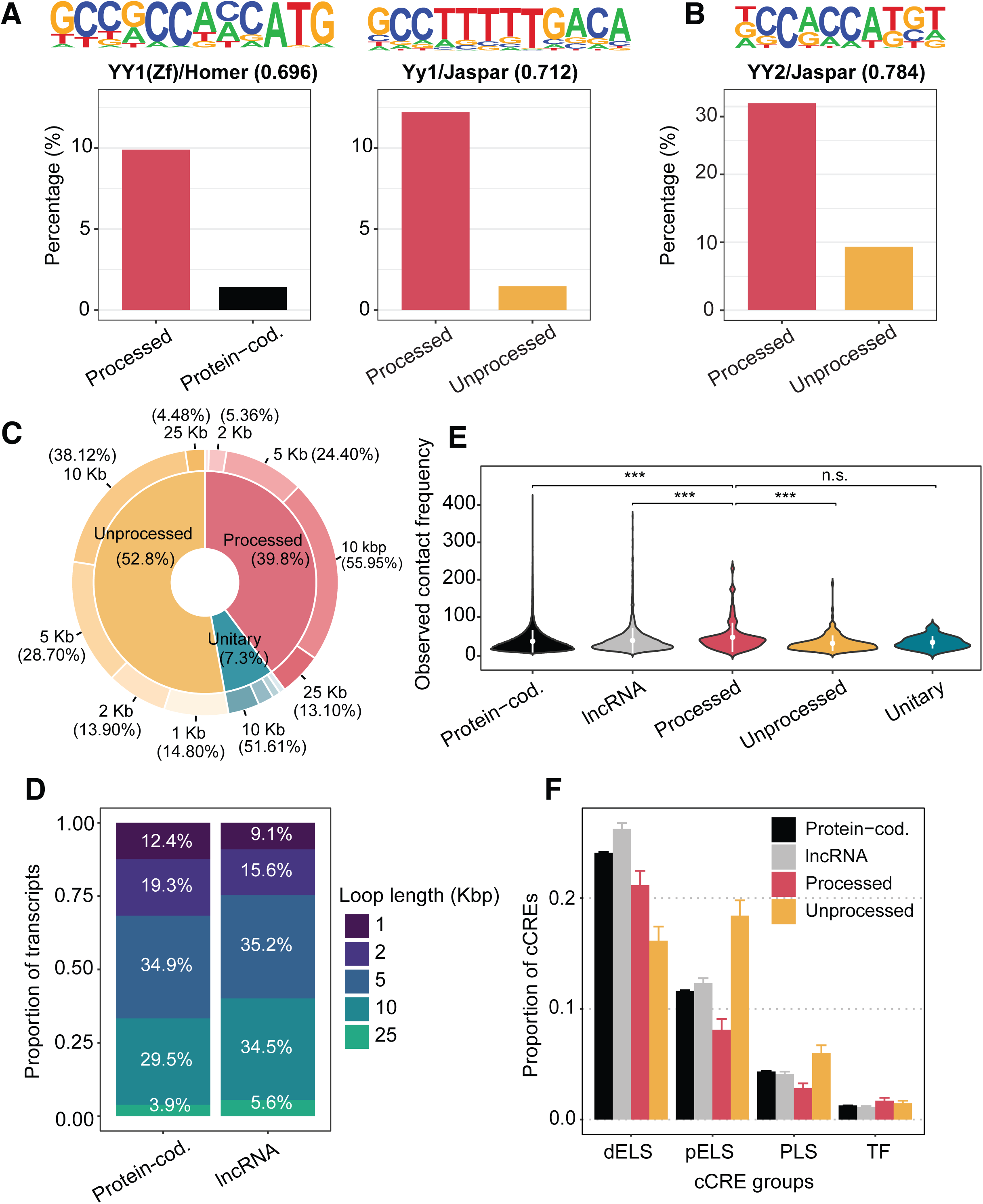
Potential factors regulating the expression of processed pseudogenes. **(A)** Bar plots displaying the percentage of promoters presenting motifs enriched in processed pseudogenes compared to promoters of protein-coding genes *(Left)* and unprocessed pseudogenes *(Right)*, respectively. Motifs are listed above the bar plots, along with their names and similarity scores compared to the best-matching ones in the database. **(B)** Similar to **(A)**, a closely related motif was identified between promoters of processed and unprocessed pseudogenes. **(C)** Pie chart showing the proportion of expressed pseudogenes from different biotypes that interact with genomic regions of varying length in the genome across tissues. **(D)** Similar to **(C),** percent stacked bar plot showing the proportion of expressed protein-coding and lncRNA transcripts that interact with genomic regions of varying lengths. **(E)** Violin plot depicting the distribution of observed contact frequency between transcripts from different gene biotypes with other genomic regions. Within each violin, the white point represents the mean, and the vertical line denotes the range of one standard deviation above and below the mean. Processed pseudogenes were used as the reference group for the two-sided Wilcoxon rank-sum test. ***, *p*-value < 0.001; n.s., not significant. **(F)** Bar plot depicting the average proportion of bases within genomic regions, where expressed transcripts from different gene biotypes interact, that overlap with the different types of annotated cCREs. Error bars represent standard errors. dELS, distal enhancer-like signature; dPLS, distal promoter-like signature; PLS, promoter-like signature; TF, transcription factor binding site. “Protein-cod.”: protein-coding.

In addition to mediating the transcription initiation of both human and mouse LINE-1s, YY1 is well-documented for its role in facilitating long-range enhancer-promoter interactions^35^. Given that processed pseudogenes lack regulatory signatures at their promoters, we explored whether they may establish contact, potentially through the involvement of YY1, with other regions in the genome that facilitate transcription initiation. We obtained Hi-C loop data from the ENCODE EN-TEx portal that matched our tissue collection and cross-referenced these data against our promoter catalog (see Methods). Across tissues, we found that pseudogenes, and in particular processed pseudogenes, tend to contact broader genomic intervals (e.g., 10 kbp and 25 kbp) compared to protein-coding and lncRNA transcripts (**Figure 4C and Figure 4D**). Furthermore, the promoters of processed pseudogenes reported a higher contact frequency than those of unprocessed pseudogenes and protein-coding transcripts (Wilcoxon rank sum test, *p*-value = 4.44×10⁻ and 6.24×10⁻, respectively, FDR-adjusted) (**Figure 4E**). To fully capture what genomic regions are specifically contacted, we intersected these intervals with annotated human candidate cis-regulatory elements (cCREs) available from the ENCODE Consortium^36^. We observed a significant difference in the regions contacted by unprocessed and processed pseudogenes. Unprocessed pseudogenes, which feature epigenetically active promoters, preferentially contacted promoter-like signature cCREs (PLS) (**Figure 4F and Figure S9**). In contrast, processed pseudogenes, which lack such active promoters, more frequently interact with distal enhancer-like signature cCREs (dELS) (**Figure 4F**). Taken together, our results suggest that the expression of processed pseudogenes might be mediated by more distal regulatory elements, with these interactions likely mediated by YY1 in a manner similar to human LINE-1.

## Discussion

In this study, we present a comprehensive catalog of promoter regions for transcribed pseudogenes, lncRNAs, and protein-coding transcripts across ∼30 human tissues, integrating high-resolution transcriptomic, epigenomic, and three-dimensional Hi-C data as well as data on genetic variation and evolutionary conservation. Collectively, our findings challenge the long-standing paradigm that pseudogenes are merely non-functional relics of evolutionary processes, demonstrating that epigenetic mechanisms, including histone modifications, nucleosome positioning, and DNA methylation, correlate with pseudogene expression, particularly for unprocessed pseudogenes. On the other hand, our results reveal an unexpected decoupling between epigenetic features and transcriptional output in processed pseudogenes. Prior to our study, it was unclear whether processed pseudogenes, following the retrotransposition event, could acquire epigenetic features resembling normal promoters and, if not, what alternative mechanisms might facilitate their transcription. Our analyses also reveal that processed pseudogene promoters exhibit distinct SNV allele frequency patterns and conservation profiles compared to unprocessed pseudogenes. Notably, we identify a potential alternative regulatory mechanism: the promoters of processed pseudogenes are enriched for YY1-binding motifs—a feature reminiscent of the human LINE-1—and frequently engage in extensive long-range chromatin interactions, preferentially contacting distal enhancer-like cCREs. These findings suggest that, in the absence of canonically active epigenetic marks, processed pseudogenes may rely on distal regulatory contacts, potentially mediated by YY1, to facilitate their transcription initiation.

Overall, we provide a refined understanding of promoter architecture and function in human tissues, which underscores the evolutionary and functional diversity of promoter regulation across distinct pseudogene subtypes, and establishes a foundation for future investigations into the regulatory roles of pseudogenes and other non-canonical transcripts.

One limitation of this study is that the analysis was based on cross-tissue multi-omics data from only four healthy individuals, which may constrain the generalizability of our findings across the broader human population. Additionally, promoters were defined based on a 2000-bp window around each TSS, which may not fully capture the complexity of promoter architecture and regulatory interactions. Ongoing efforts by the GENCODE project, including the integration of CapTrap-Seq data^37^ and the application of the deep learning model ProCapNet^38^, aim to refine promoter annotations and improve accuracy in defining TSSs. Furthermore, our analysis focused primarily on canonically active histone marks to assess their patterns across transcribed pseudogenes. Future studies may investigate the potential role of repressive histone modifications, such as H3K9me3 and H3K27me3, in regulating pseudogene expression, particularly in the context of promoter silencing and heterochromatin-associated repression.

Given the increasing evidence implicating pseudogenes in human development^39^ and various diseases^40–42^, further exploration of their epigenetic and transcriptional patterns may reveal novel biological roles and potential therapeutic targets. As our understanding of the non-coding genome continues to evolve, pseudogenes should no longer be regarded as merely vestigial elements but rather as epigenetically controlled genomic entities with potentially functional significance. We anticipate that the catalog and integrative analyses developed in this study will serve as a resource for the research community, fostering continued exploration into the regulatory potential of the non-coding genome.

## Methods

### Assigning promoter regions of pseudogene, protein-coding, and lncRNA transcripts

To streamline the analysis of numerous functional assays and maintain consistency with the ENCODE uniform analysis pipelines^43^, in this study, we utilized the GENCODE v29 annotation based on the GRCh38 human genome assembly. We included genes and their associated transcripts, focusing specifically on the following gene biotypes:

1. Protein-coding genes: indicated by “protein_coding”.
2. Long non-coding RNA (lncRNA) genes: encompassing “3prime_overlapping_ncrna”, “antisense”, “’bidirectional_promoter_lncrna”, “macro_lncRNA”, “non_coding”, “processed_transcript”, “sense_intronic”, “’sense_overlapping”, and “lincRNA”.
3. Pseudogenes: encompassing “processed_pseudogene”, “unprocessed_pseudogene”, “unitary_pseudogene”, “transcribed_processed_pseudogene”, “transcribed_unprocessed_pseudogene”, “transcribed_unitary_pseudogene”, “translated_processed_pseudogene”, “translated_unprocessed_pseudogene”.

For these transcripts, promotors were defined as ± 1000 bp windows upstream and downstream of the TSSs.

### Filtering transcripts by annotation, expression, and RAMPAGE peaks

To unambiguously characterize promoters, we removed transcripts whose promoters overlap with those of transcripts from different genes in a head-to-head, tail-to-tail, or unidirectional manner^44^ in subsequent analyses. In other words, in a unidirectional overlap, both promoters are located on the same strand, where one starts before the other has ended, resulting in an overlap. In head-to-head and tail-to-tail overlaps, the promoters are on opposite strands. Head-to-head indicates that the starting sites face each other, whereas tail-to-tail means the termination sites are aligned. In such a way, we have included in a total of 14,253 pseudogenes, 17,624 lncRNAs, and 106,643 protein-coding transcripts.

We downloaded isoform-level quantifications from the ENCODE EN-TEx portal (http://entex.encodeproject.org/) for matched human tissues from 4 donors. Transcripts with an average expression level below 1 TPM across donors and technical replicates were excluded. Finally, for each gene and tissue, the most abundantly expressed transcript was selected as its representative.

RAMPAGE was capable of capturing 5’-complete complementary DNAs (cDNAs), allowing to precisely identify TSSs and accurately quantify promoter activity^45^. To validate our assignment of promoters, we leveraged an integrated collection of representative RAMPAGE peaks^17^. We employed BEDTools intersect^46^ to find potential overlaps between the promoters of expressed transcripts and representative RAMPAGE peaks. If a RAMPAGE peak overlaps with a promoter by at least 1 bp, we consider the promoter to be experimentally supported.

### Profiling epigenetic patterns with matched tissue-specific data

To characterize the epigenetic patterns associated with each transcript, we downloaded tissue-specific histone ChIP-Seq peaks, ATAC-Seq peaks, and DNase-Seq peaks from the ENCODE EN-TEx portal (http://entex.encodeproject.org/) across multiple human tissues from 4 donors. Specifically, we downloaded six histone ChIP-Seq peaks for H3K27ac, H3K4me3, H3K4me1, H3K36me3, H3K9me3, and H3K27me3. Certain tissues, such as the omental fat pad and subcutaneous adipose tissue, were excluded due to insufficient RNA-Seq and/or histone ChIP-Seq data. Thus, we have tissue-specific epigenomic data for 26 human tissues. For each assay and tissue, we merged peaks across donors and technical replicates using BEDTools merge^46^ with the -i stdin - c 1 -o count options. Then, we employed BEDTools intersect^46^ with the -wao option to calculate the base-pair overlap between defined promoters and peaks derived from histone ChIP-Seq, ATAC-Seq, and DNase-Seq data (**Table S1**). For those peaks, we calculated the proportion of bases (i.e., coverage) overlapping peaks for each transcript in each tissue due to the tissue-specific nature of these signals. We then aggregate them, calculating the average proportion of coverage across transcripts and tissues for each biotype.

Additionally, we obtained tissue-specific DNA methylation profiling by array assay across the same tissues. To ensure consistency across donors and technical replicates for each tissue, we used BEDTools merge^46^ with the -c 5 -o mean options to calculate the average beta value for each CpG site. We classified CpG sites based on their beta values into three categories: hypomethylated (beta value < 0.2), hypermethylated (beta value > 0.8), and heterogeneously methylated (0.2 ≤ beta value ≤ 0.8)^47^. For each transcript, we quantified the number of CpG sites based on probe detection. We then categorized the CpG sites according to their methylation status and calculated the proportion of each category across tissues.

We further utilized *in situ* HiC data available for two tissues, the transverse colon and skeletal muscle, to support our findings. Genome compartment files of each tissue were downloaded and merged across donors using wiggletools mean^48^ separately. Next, we filtered transcripts specifically expressed in these two tissues and employed bigWigAverageOverBed^49^ to compute the average signal intensity over the respective promoters of these transcripts.

### Assessing inter- and intra-species conservation of selected regions

Given that most processed pseudogenes in the human genome originated during a burst of retrotransposition events at the dawn of the primate lineage^50^, we obtained 20-way phastCons scores from the UCSC Genome Browser, including alignments with primates, mouse, and dog (https://hgdownload.cse.ucsc.edu/goldenpath/hg38/phastCons20way/) to assess inter-species conservation. We then used bigwigAverageOverBed^49^ to calculate conservation scores separately for the upstream and downstream regions of the TSSs, as well as for the exonic regions of the transcripts. For a given transcript, its conservation score is determined by calculating the weighted average conservation across its exonic regions.

To assess intra-species conservation, we obtained the genome-wide SNV data from gnomAD v4 (https://gnomad.broadinstitute.org/data#v4-variants/)^16^. We then used VCFtools^51^ with the --remove-filtered-all --min-alleles 2 --max-alleles 2 options to retain high-quality variants and restrict the dataset to biallelic sites. Given the substantial allele frequency differences among populations^52^, we designated the non-Finnish European (NFE, N = ∼600K) group as the reference population and excluded sites with an MAF below 1x10^-5^ from our analysis. We used BEDTools intersect^46^ to identify all variants located in the upstream and downstream regions of the TSSs, as well as within the exonic regions of the transcripts. The MAF and Phred-scaled CADD score for each SNV were directly retrieved from the dataset. To calculate the proportion of ultra-rare variants within a given region, we classified SNVs with an MAF below 1×10⁻ as ultra-rare, given the large size of our cohort.

### Mapping the connection between transposable elements and processed pseudogenes

To investigate the potential shared regulatory mechanisms between processed pseudogenes and TEs, we obtained genome-wide TE annotation in the GRCh38 genome assembly from RepeatMasker open-4.0.5 (https://www.repeatmasker.org/species/hg.html). We analyzed promoters of expressed transcripts across multiple tissues by intersecting their genomic coordinates with annotated TEs and quantified the extent of overlap based on the number of overlapping base lengths. For each promoter, we cataloged the diversity of intersecting TE classes, including long-terminal repeats (LTR), DNA transposons (DNA), simple repeats, low complexity, SINEs, and LINEs. To assess the similarity between TEs and promoters, we calculated the Jaccard index, defined as the ratio of the length of the overlapping region to the total combined length of the TE and promoter. This metric provided a normalized measure of the extent of overlap.

### Identifying enriched motifs within promoter regions

Motif enrichment analysis was performed using Homer2^29^ with the findMotifsGenome.pl script, employing the options -size 200 -S 10 -bg. Promoter regions of processed pseudogenes were selected across tissues and compared to those of unprocessed pseudogenes and protein-coding transcripts to identify enriched known and *de novo* motifs. For *de novo* motifs, the similarity score between each identified motif and its closest match in the reference database (e.g. Homer and Jaspar) was reported.

### Identifying genomic regions contacted by processed pseudogenes by Hi-C data

We obtained intact Hi-C loop data from the ENCODE portal as part of the ENCODE Phase 4 project. For transcripts expressed in each tissue, we used BEDTools pairtobed^46^ with the -type either option to identify overlaps between these transcripts and tissue-specific HiC loop regions. Then, we aggregated these regions that interacted with transcripts across all tissues and intersected them with the catalog of annotated human cCREs v4 obtained from SCREEN (https://screen.wenglab.org/).

### Quantification and statistical analysis

All analyses and statistical tests in this study were conducted using Python (version 3.9.13) and R (version 4.2.0), as detailed in the Methods and figure legends. Unless otherwise specified, plots were made with the ggplot2^53^ package in R. All box plots depict the first and third quartiles as the lower and upper bounds of the box, with a central band showing the median value and whiskers representing 1.5x the interquartile range (IQR).

## Data and code availability

No new data were generated for this study. All datasets used are publicly available and can be accessed from the ENCODE EN-TEx portal (http://entex.encodeproject.org/) and other sources specified in the Methods section. All custom scripts and some intermediate files in this study are available on GitHub (https://github.com/gersteinlab/epiPgene).

## Author contributions

YJ, BB, and MG conceived the project and designed the study. YJ performed the computational analyses. YJ and BB wrote the original draft of the manuscript, which MG subsequently revised.

## Competing interests

The authors declare no competing interests.

## Notes

### Competing Interest Statement

The authors have declared no competing interest.

