## Supplementary Information for "Epigenetic characterization of pseudogenes across human tissues"

Supplementary Figures 1-9

Supplementary Tables 1-2

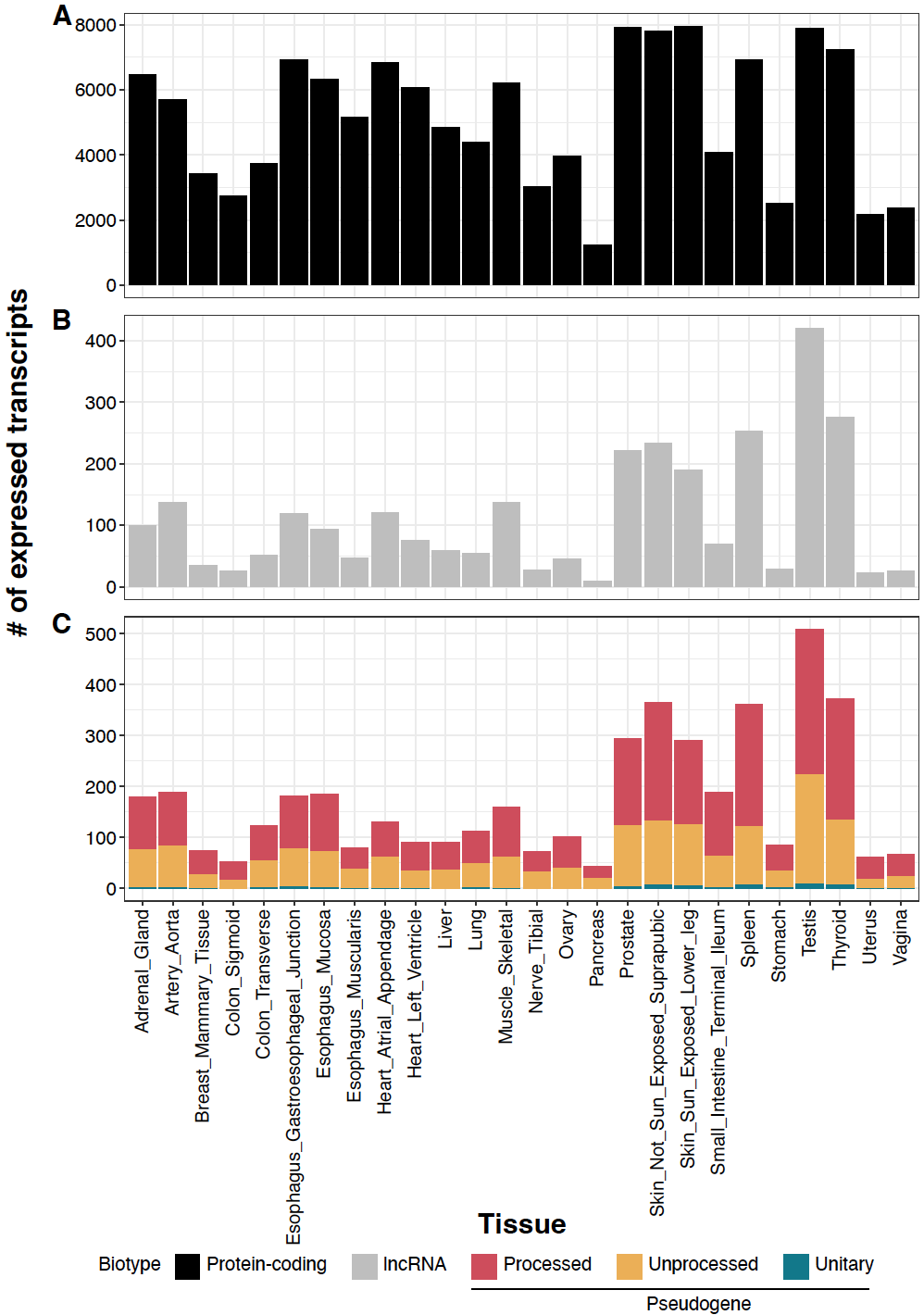

**Figure S1.** The number of expressed transcripts in GENCODE v29 across various tissues for **(A)** protein-coding transcripts, **(B)** lncRNAs, and **(C)** pseudogenes, further categorized into processed, unprocessed, and unitary pseudogenes.

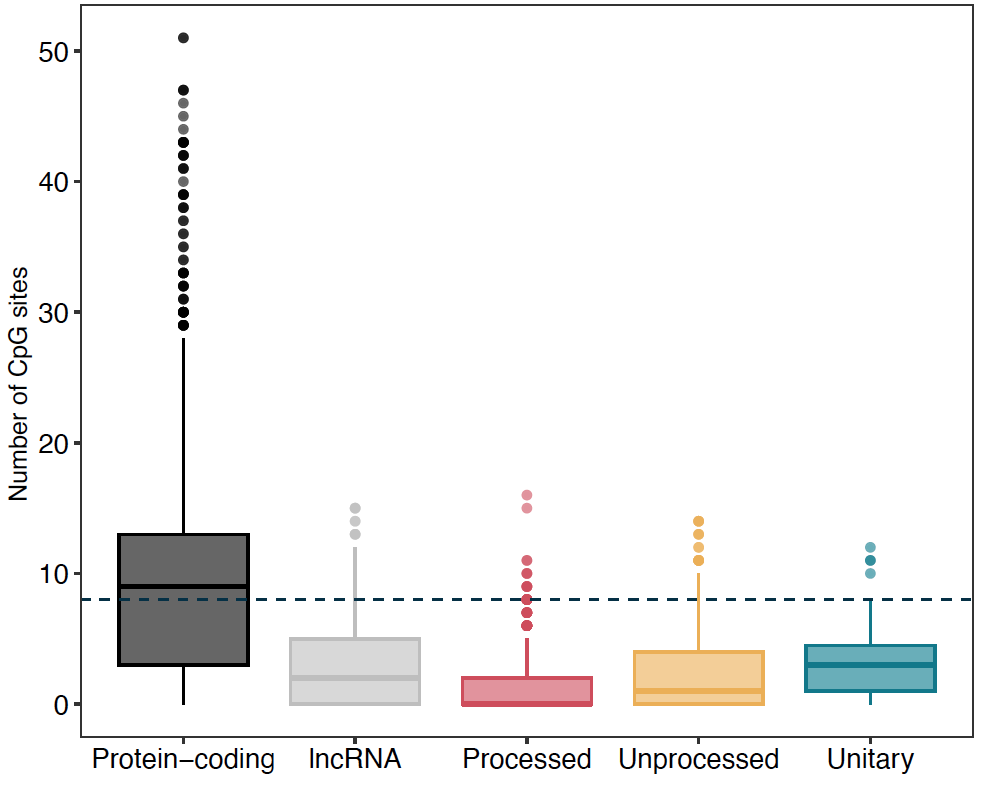

**Figure S2.** The number of CpG sites within promoters of expressed transcripts from different gene biotypes. Horizontal lines represent the median number of CpG sites across all transcripts.

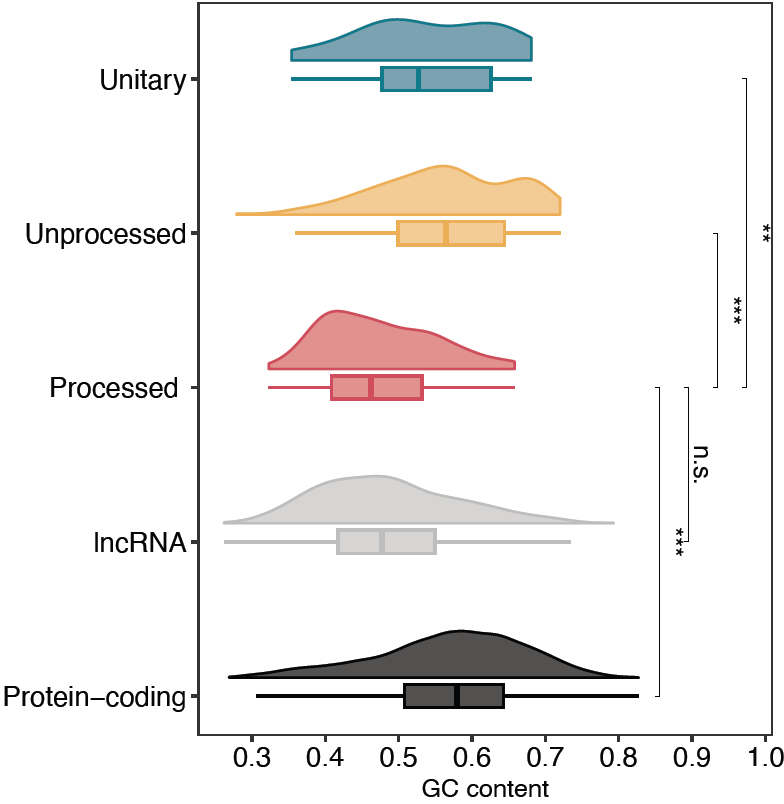

**Figure S3.** The GC content of regions downstream of TSS across transcripts from different gene biotypes. Processed pseudogenes were used as the reference group for the two-sided Wilcoxon rank sum test. ***, *p*-value < 0.001; **, *p*-value < 0.01; n.s., not significant.

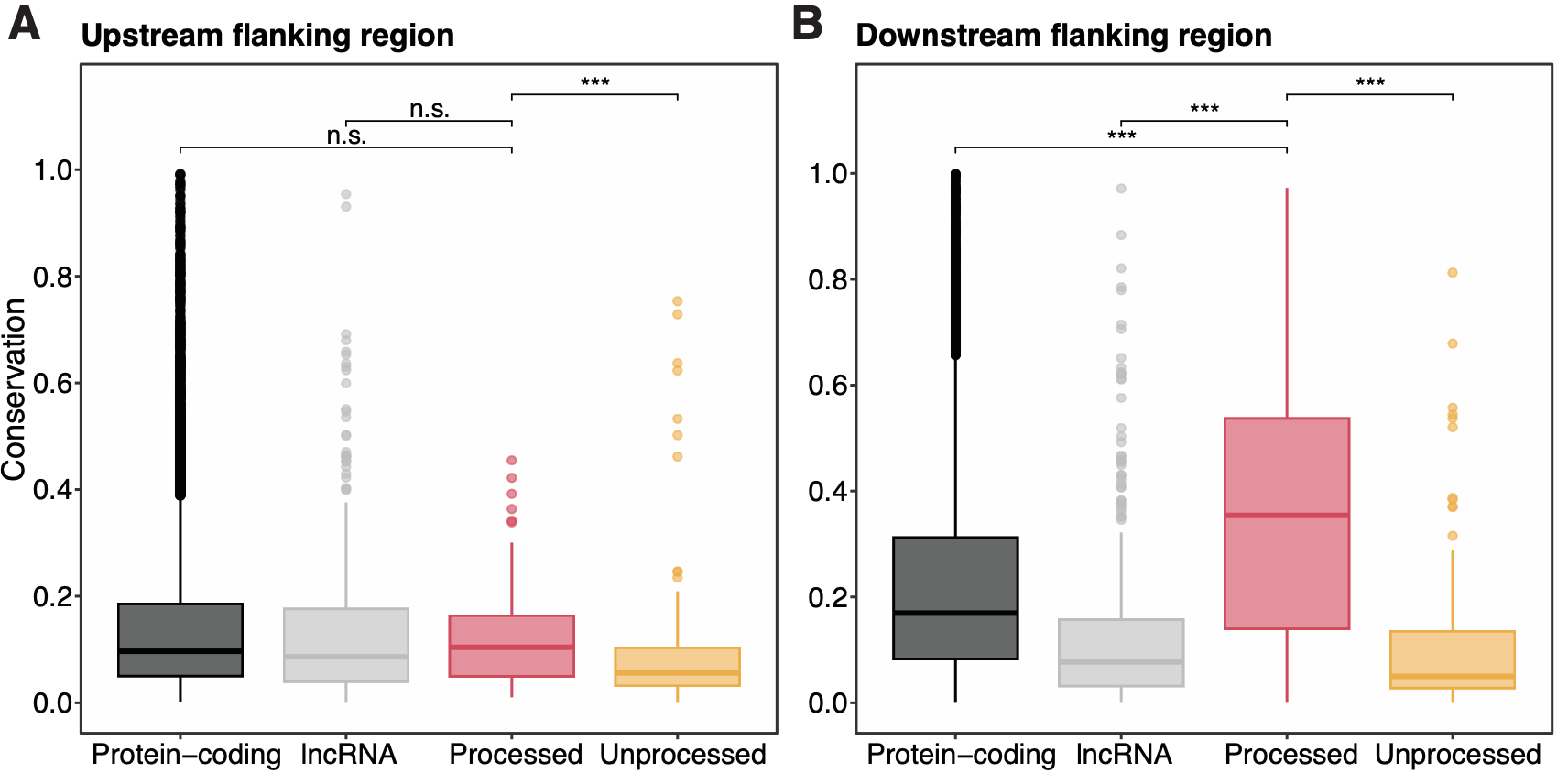

**Figure S4.** 20-way phastCons conservation scores genomic regions flanking TSS: **(A)** upstream and **(B)** downstream regions. Processed pseudogenes were used as the reference group for the two-sided Wilcoxon rank sum test. ***, *p*-value < 0.001; n.s., not significant.

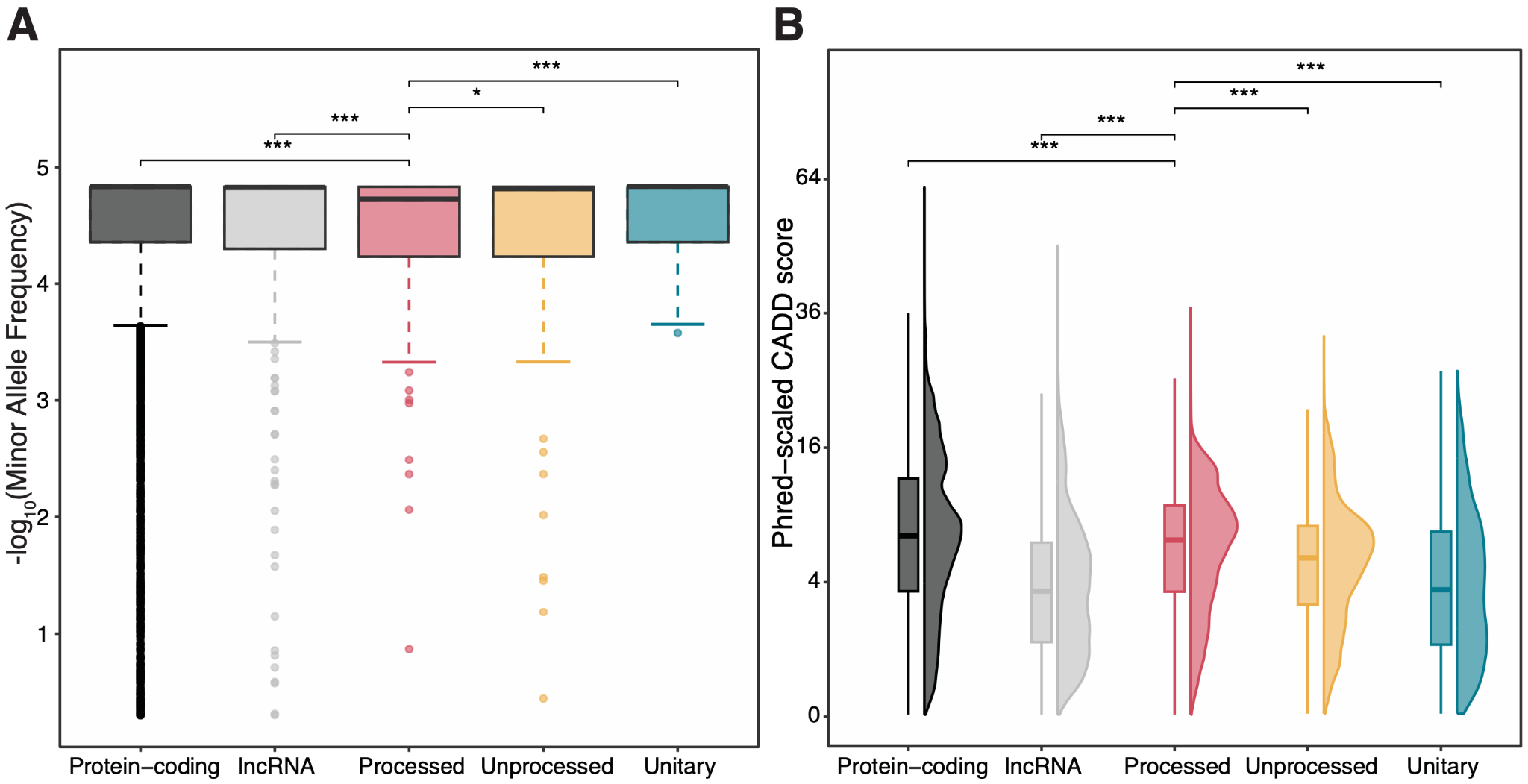

**Figure S5.** The distribution of MAF **(A)** and Phred-scaled CADD scores **(B)** for SNVs within the downstream regions of TSS. Due to the large sample size, only 0.5% of SNVs classified as outliers were randomly selected for visualization in **(A)**. Processed pseudogenes were used as the reference group for the two-sided Wilcoxon rank sum test. ***, *p*-value < 0.001; *, *p*-value < 0.05.

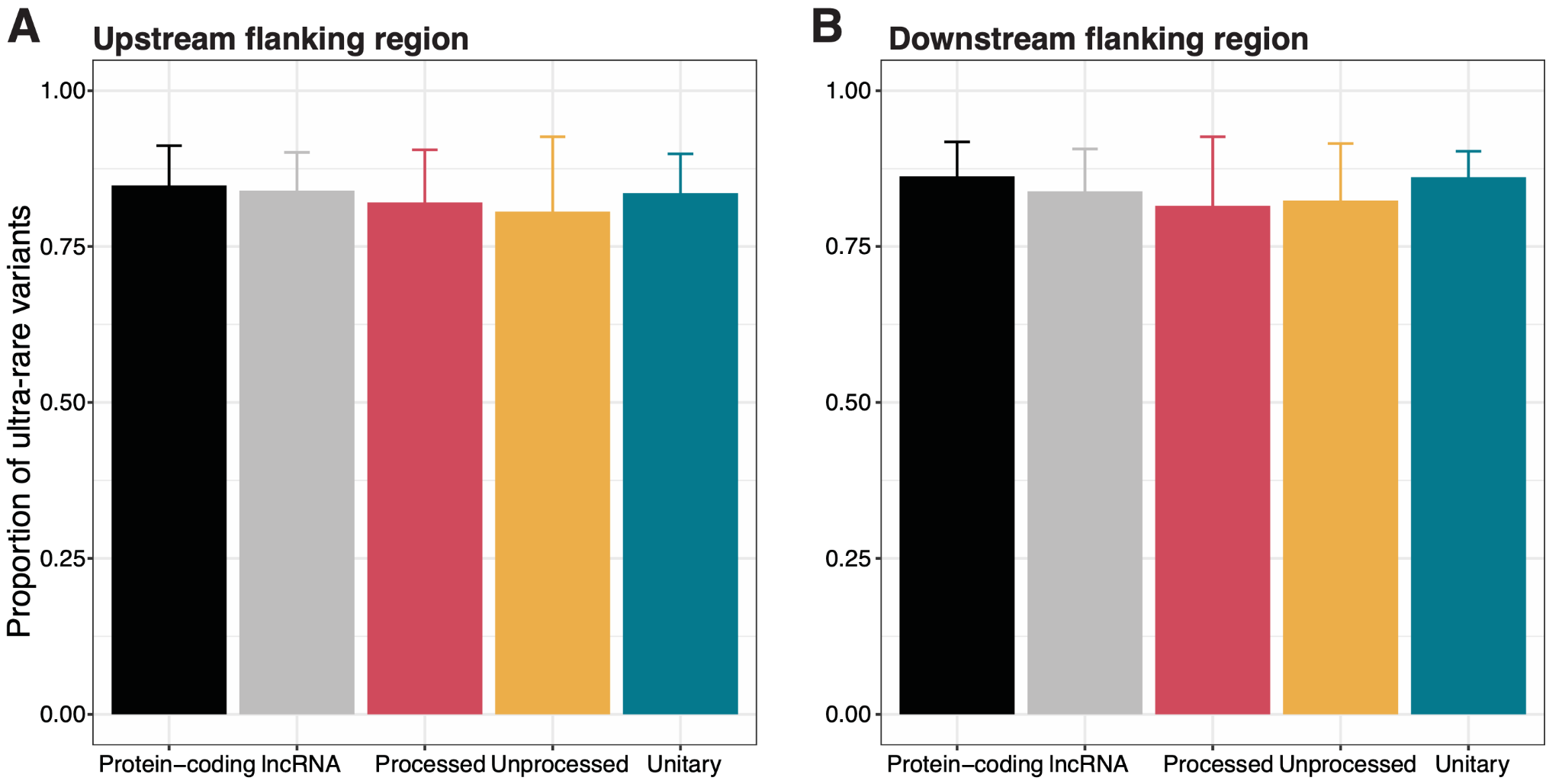

**Figure S6.** The average proportion of ultra-rare variants in the upstream **(A)** and downstream **(B)** regions of TSS. Error bars represent standard deviation.

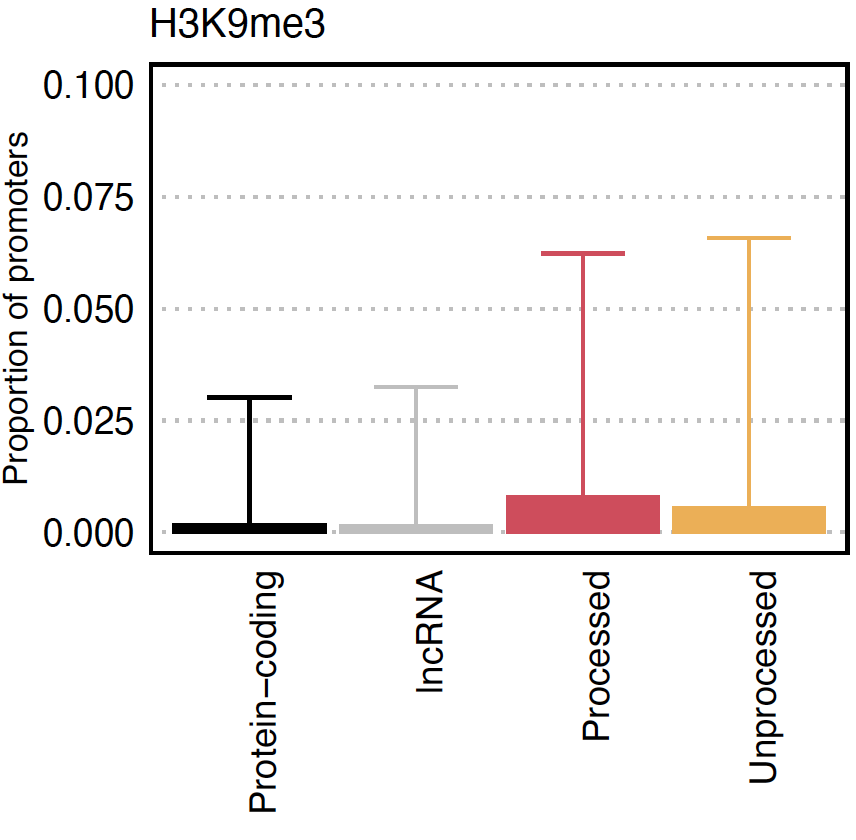

**Figure S7.** The average proportion of bases in promoters of protein-coding, lncRNA, processed pseudogenes, and unprocessed pseudogenes covered by peaks from the repressive mark, H3K9me3. Proportions were calculated for each transcript in each tissue and then aggregated across tissues for each gene biotype. Error bars represent standard deviations across tissues.

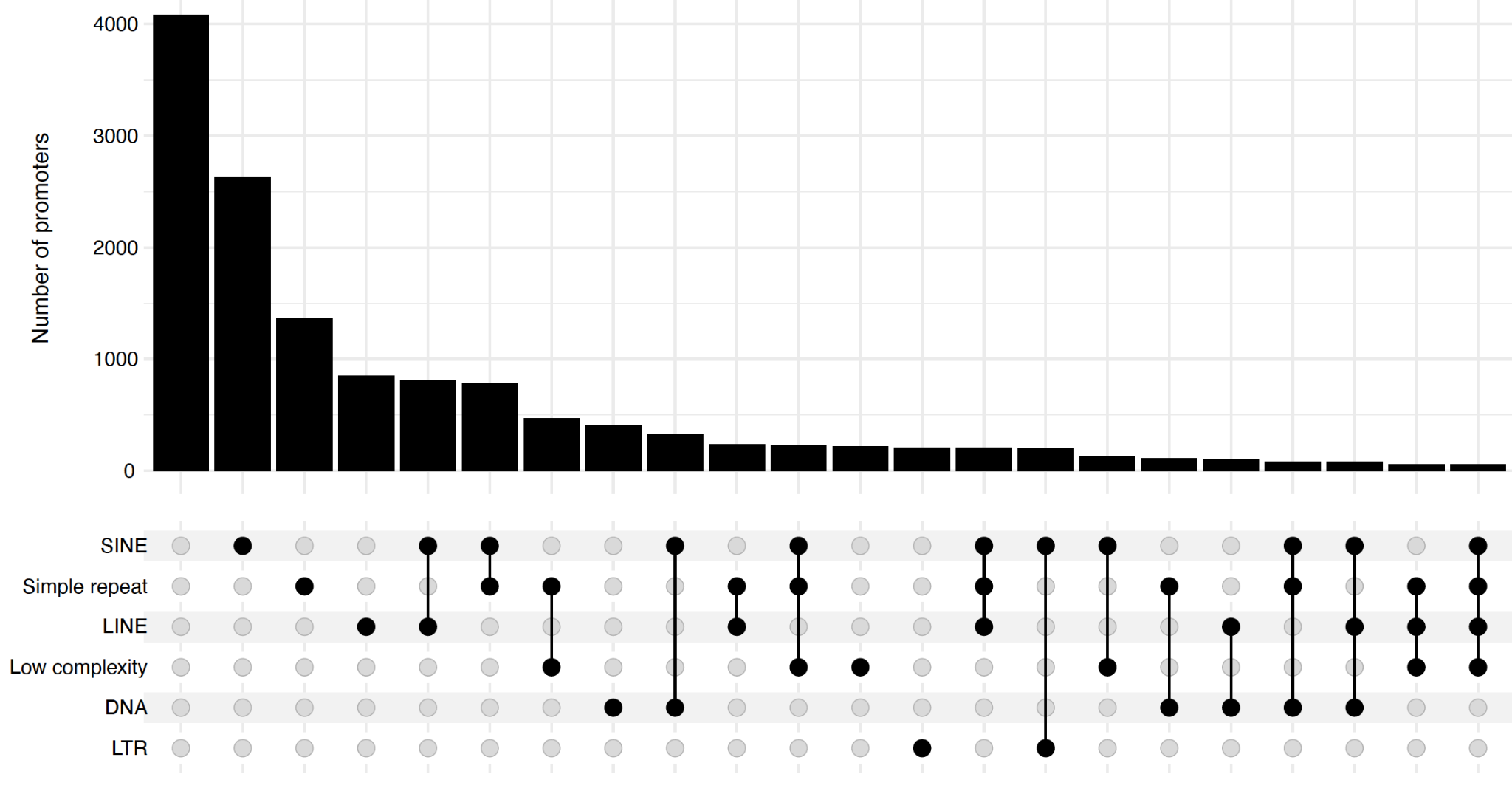

**Figure S8.** Upset plot for promoters of protein-coding transcripts overlapping with various transposable elements. Some intersection groups were excluded due to the small size of elements (N < 50).

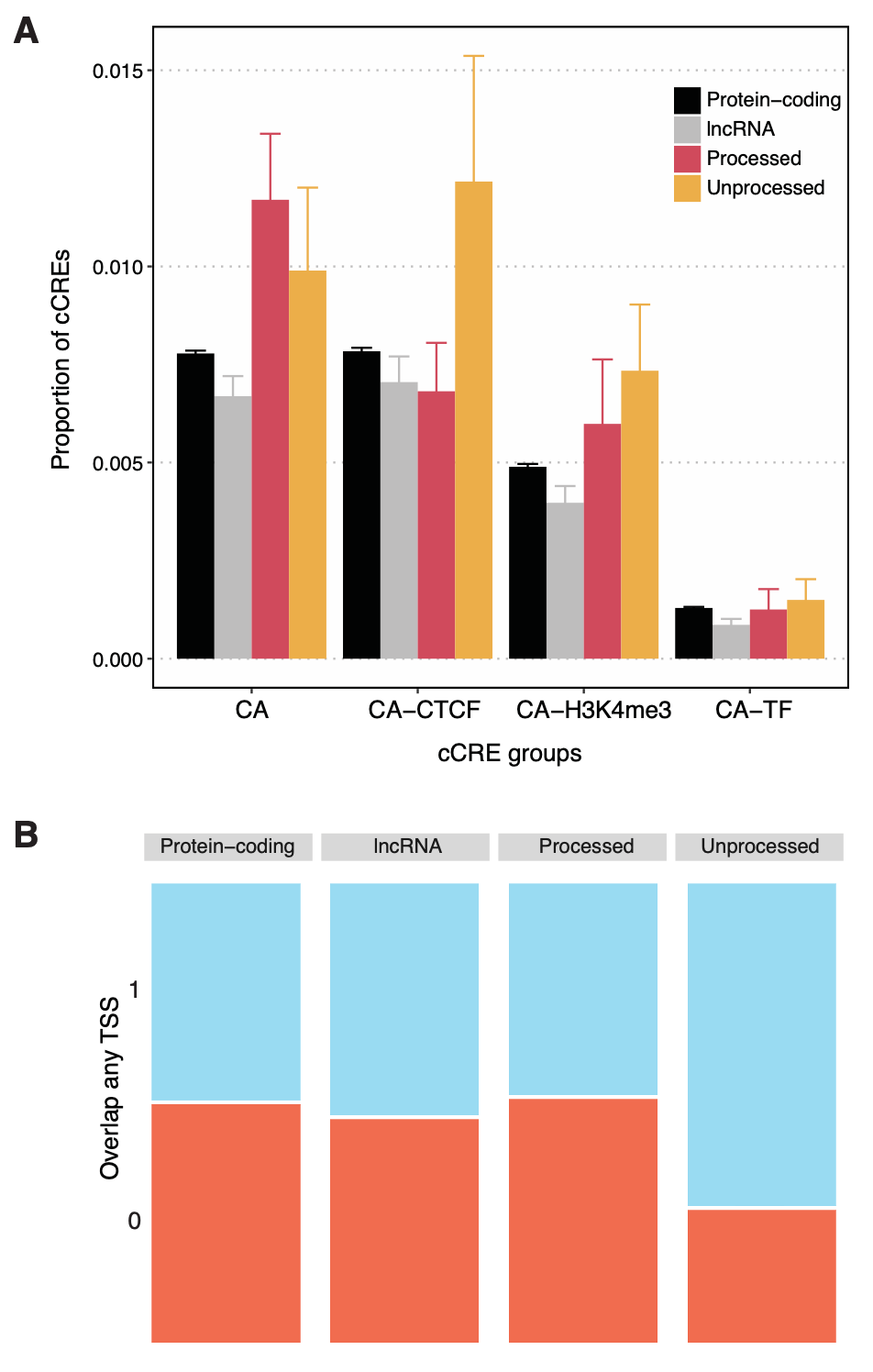

**Figure S9.** The proportion of bases within genomic regions, where expressed transcripts from different gene biotypes interact, that overlap with the different types of annotated cCREs associated with chromatin accessibility (CA) (A) and annotated TSSs (B) in GENCODE v29 of the human genome. Overlap status is indicated as 1 (Yes) or 0 (No). Error bars represent standard errors.

**Table S1.** Comprehensive annotation of promoter activity across human tissues. The table is available via Zenodo (<https://doi.org/10.5281/zenodo.14934024>). The dataset contains the following columns:

- 1-3. BED intervals that define the promoter regions (TSS ± 1 kbp).
- 4. Ensembl gene ID corresponding to the associated transcript.
- 7. Ensembl transcript ID to which the promoter was assigned.
- 8. Gene biotype, as annotated in GENCODE v29.
- 9. Tissue name.
- 10. Average transcript-level expression (in TPM) across donors and technical replicates in the tissue.
- 11–16. Length of overlap between the promoter regions and peaks from various histone mark ChIP-seq datasets, including H3K27ac, H3K4me3, H3K27me3, H3K36me3, H3K4me1, and H3K9me3. "NA" indicates missing experimental data, and a value of "0" indicates no overlap between the promoter region and the corresponding peak.
- 17–18. Length of overlap between the promoter regions and peaks from DNase-Seq and ATAC-Seq data.
- 19-22. Number of CpG sites within the promoters and different levels of methylation.

The

**Table S2.** Functional properties of upstream regions of TSS. The table is available via Zenodo (<https://doi.org/10.5281/zenodo.14934024>). The dataset contains the following columns:

- 1-3. BED intervals that define regions upstream of TSS.
- 4. Ensembl gene ID corresponding to the associated transcript.
- 7. Ensembl transcript ID to which the promoter was assigned.
- 8. Gene biotype, as annotated in GENCODE v29.
- 9. GC content of the selected region
- 10. 20-way phastCons conservation score of the selected region
- 11-15. The number of genetic variants within the selected region, along with the proportion of ultra-rare (MAF < 1x10^-4^), rare (MAF < 1x10^-3^), low-frequency (MAF < 0.05), and common variants.
- 16-18. First, second, and third quantiles of phred-scaled CADD scores for variants within the selected region.
